# A pair of primers facing at the double-strand break site enables to detect NHEJ-mediated indel mutations at a 1-bp resolution

**DOI:** 10.1101/2022.02.07.479376

**Authors:** Faryal Ijaz, Ryota Nakazato, Mitsutoshi Setou, Koji Ikegami

**Affiliations:** Department of Anatomy and Developmental Biology, Graduate School of Biomedical and Health Sciences, Hiroshima University, 1-2-3 Kasumi, Minami-Ku, Hiroshima 734-8553, Japan; Department of Cellular and Molecular Anatomy and International Mass Imaging Center, Hamamatsu University School of Medicine, Hamamatsu 431-3192, Japan; JST, PRESTO, 4-1-8 Honcho, Kawaguchi, Saitama 332-0012, Japan

## Abstract

The introduction of small insertion/deletion (indel) mutations in the coding region of genes by the site-specific nucleases such as Cas9 allows researchers to obtain frameshift null mutants. Technically simple and costly reasonable genotyping methods are awaited to efficiently screen the frameshift null mutant candidates. Here, we developed a simple genotyping method called DST-PCR (Double-strand break Site-Targeted PCR) using “face-to-face” primers where the 3’ ends of forward and reverse primers face each other at the position between 3-bp and 4-bp upstream of the PAM sequence, which is generally the Cas9-mediated double-strand break site. Generated amplicons are directly subjected to TBE-High-Resolution PAGE, which contains a high concentration of bis-acrylamide, for mutant clones detection with 1-bp resolution. We demonstrate six examples of CRISPR/Cas9-engineered knockout cells, to screen indels to obtain potential KO cell clones utilizing our approach. This method allowed us to detect 1-bp to 2-bp insertion and 1-bp to 4-bp deletion in one or both alleles of mutant cell clones. In addition, this technique also allowed the identification of heterozygous and homozygous biallelic functional KO candidates. Thus, DST-PCR coupled with TBE-High-Resolution PAGE is a simple method for genotyping of the CRISPR/Cas9-engineered cell lines, which can be performed without special equipment and techniques.

## Introduction

A prokaryotic RNA-mediated adaptive immune system known as clustered regularly interspaced short palindromic repeats (CRISPR) / CRISPR-associated protein 9 (Cas9) system has been developed into a revolutionary genome-editing technology [1]. The Cas9 nuclease combined with a short single-guide RNA (sgRNA) allows researchers to create site-specific double-strand breaks (DSBs) [2–4]. These DSBs are predominantly repaired by error-prone non-homologous end-joining (NHEJ) pathway often producing mutations of small nucleotide substitutions or insertion/deletion (indel) at the targeted sequence [5]. These indels are of interest to researchers to acquire frameshift null mutants for gene functional studies [6].

One of the major challenges to perform screening of knockout (KO) clones that possess CRISPR/Cas9-induced frameshift mutations is the need for simple and cost-effective strategies that do not require special equipment and are available in standard laboratories and institutes. Western blot analyses can be chosen for screening if reliable antibodies are available and the samples are cell lines. However, in many cases, genotyping with genomic DNA is required in cases where reliable antibodies are not available or the samples are not cell lines, e.g. animals. In these cases, initial screening is important to choose “best possible” candidates before sequencing.

The T7E1/Surveyor assays [7, 8] are broadly used for detecting modified genes. They can detect heteroduplexes formed by the hybridization of wild-type and mutated DNA strands or two differently mutated DNA strands [9]. A weak point of the technique is that it cannot differentiate homozygous biallelic mutants from wild-type nor heterozygous biallelic mutants from heterozygous monoallelic mutants. In addition, this technique does not provide information about types of mutations as well as the number of indels, which hampers researchers to screen out indels of nucleotides of multiples of three. Another commonly used strategy for genotyping is CRISPR/Cas9-derived RNA-guided engineered endonucleases in Restriction fragment length polymorphism analysis (RGEN-RFLP) analysis [10]. Unlike T7E1/Surveyor assay, it can detect homozygous biallelic mutants. This assay, however, still cannot provide information about the presence or absence of frameshift mutations.

Heteroduplex Mobility Assay (HMA) can also distinguish between the heteroduplexes and homoduplexes DNA in native PAGE [11,12]. Moreover, improved versions of the HMA called improved heteroduplex analysis (iHDA) [13] and Probe-Induced HMA (PRIMA) [14] use DNA probes to detect 1-bp pair insertions after gene editing. A weak point of these methods is that they require special equipment and might need skilled techniques for optimizations.

In this paper, we present an alternative simple one-step PCR-based method, namely <u>D</u>SB <u>S</u>ite-<u>T</u>argeted PCR (DST-PCR) coupled with Tris-borate-EDTA high concentration bis-acrylamide gel electrophoresis (TBE-PAGE) for screening indels introduced by CRISPR/Cas9 genome editing in cultured cells. DST-PCR approach demonstrated here is a simple PCR with neither any special techniques nor special equipment and enables genotyping with the 1-bp resolution. We here show examples of genotyping for six different genes.

## Materials and Methods

### Cell culture

NIH/3T3 cells, a fibroblast cell line derived from the whole murine embryo, were purchased from American Type Culture Collection (ATCC CRL-1658). IMCD3 cells, inner medullary collecting duct cell line derived from mouse, were purchased from American Type Culture Collection (ATCC CRL-2123). NIH/3T3 cells were cultured in DMEM-High glucose (044-29765, Wako, Japan) containing 10% fetal bovine serum (FBS) (26140079, Gibco). IMCD3 were cultured in DMEM / Ham’s F-12 (048-29785, Wako, Japan) containing 10% FBS. All cell lines were maintained in a humidified incubator at 37°C supplied with 5% CO_2_ in the air. Cells were induced for ciliogenesis with the following conditions: Confluent NIH/3T3 cells were cultured in DMEM-High Glucose with 1% FBS for 12 h. Confluent mIMCD-3 were cultured in serum-free DMEM/ Ham’s F-12 medium for 12 h.

### Construct design

To construct all-in-one expression plasmids for knock-out, a 20-bp target sequence was sub-cloned into U6-gRNA/CMV-Cas9-2A-GFP plasmid backbone (ATUM, CA, USA). The target sequences of guide RNA were selected using Broad Institute GPP sgRNA Designer (https://portals.broadinstitute.org/gpp/public/analysis-tools/sgrna-design). Target sequences used in this study are listed as follows.

mDync2h1: TATACATACGAGTACCAGGT;
mIft144: TAAGGATAATCTAACCAGTG;
mInpp5e:AGTGATCGTCACCAGCCAAG;
mArntl: TTGTCGTAGGATGTGACCGA;
mPkd1: CATCATGCTGTAAGCCAATG;
mPkd2: CCAATGTGTACTACTACACT;

### Generation of knockout cell lines

Knockout cell lines were generated with a CRISPR/CAS9-based genome editing technique as described in previous studies [15, 16]. Cells were plated at a density of 2 x 10^4^ per cm^2^ and were transfected with all-in-one plasmid U6-gRNA/CMV-Cas9-2A-GFP using polyethylenimine (PEI) [17]. After 3 days of transfection, GFP positive cells were sorted using a FACS Aria2 SORP cell sorter (Becton Dickinson) and were plated in an appropriate culture medium containing 1% Penicillin/Streptomycin (168-23191, Wako, Japan) in a 12-well plate. The cells were later expanded into a 10-cm dish. Then, the cells were collected and single cell-cloned into 96-well plates with the limiting dilution-culture method at 1 cell/well. Cells were allowed to grow for 1-2 weeks. For the selection of monocolonies, microscopic observations were made to monitor single-cell colony formation and confluency. Selected colonies were expanded into the duplicate of multiwell plates from which one culture was used to make frozen stocks for subsequent use and the other culture for screening by genotyping. To extract genomic DNA cells were first lysed in lysis buffer (50mM Tris, pH 7.5, 100 mM EDTA, 1% SDS) containing Proteinase K (0.4 mg/mL) at 70°C for 60 min. The volume of the lysis buffer was adjusted according to the confluency of cell clones in each well. Half a volume of absolute ethanol was added to the mixture. Then, genomic DNA was isolated using Favor Prep Blood Genomic DNA Extraction Mini Kit (FABGK001-2, Favorgen Biotech Corp) and used for screening of mutant clones by DST-PCR.

### Genotyping of CRISPR mutants with DST-PCR

Primers were designed based on the following criteria: the 3’ end of both forward and reverse primers faced each other at the position between 3-bp and 4-bp upstream of the PAM sequence. The primers were designed for a product size ranging from 39-to 41-bp. Smaller amplicons (<40-bp) are preferable to detect 1-bp indels. The list of primers used for genotyping and estimated amplicon size are listed in Table 1. The PCR fragment was prepared with a normal PCR protocol using KOD-Plus-Neo DNA polymerase (KOD-401, Toyobo, Japan) or ExTaq DNA Polymerase (RR001, Takara, Japan), both of which had 3’-5’ exonuclease activities.

**Table 1.**
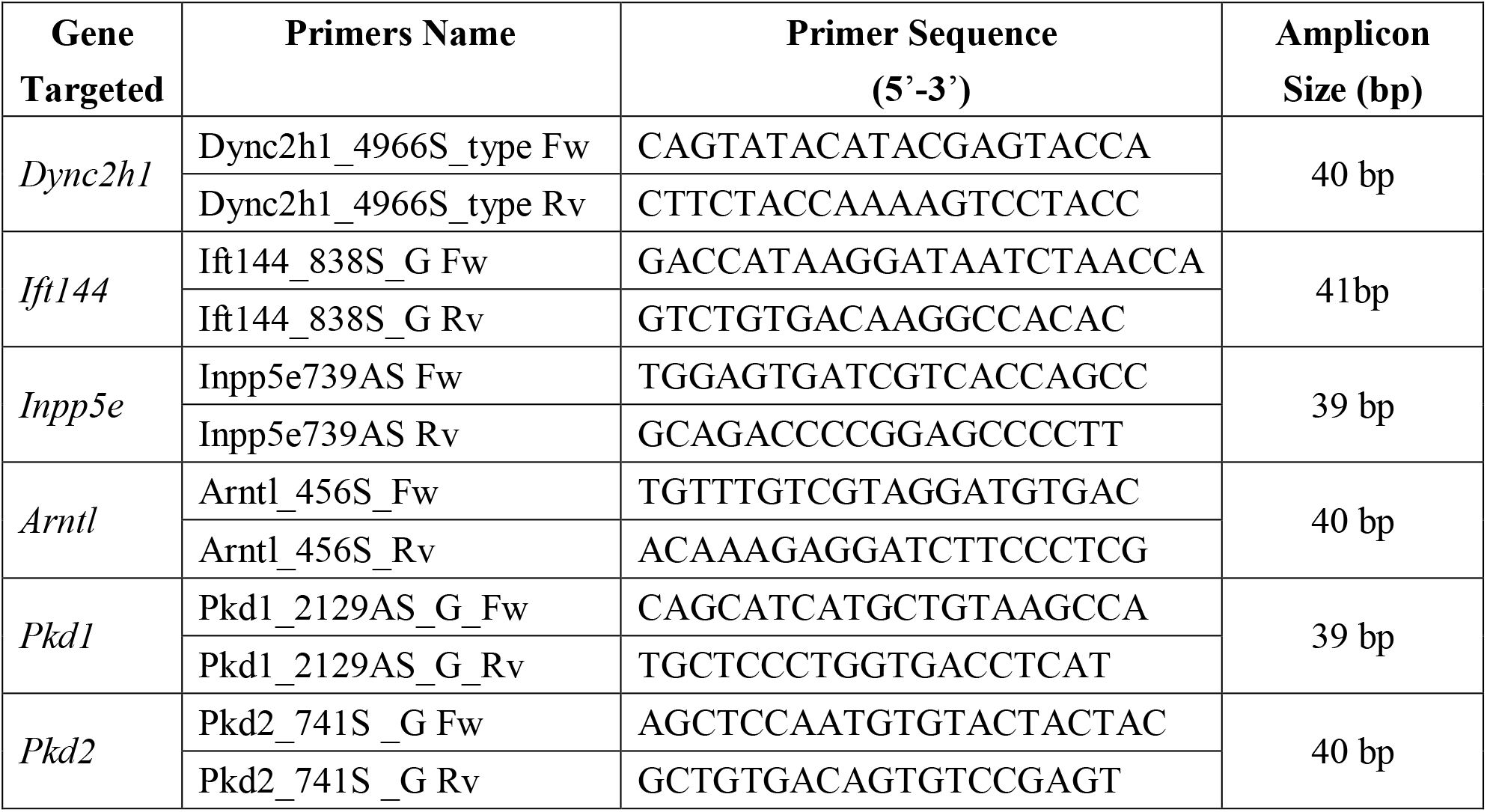
DST-PCR primers for screening.

Twelve to 15% polyacrylamide gels were prepared using a 40% (w/v) Acrylamide/Bis mixed solution containing 38% (w/v) Acrylamide and 2% (w/v) bis-Acrylamide for a monomer to crosslinker ratio of 19:1 (Wako, Japan, 013-25675) in 1X Tris-Borate-EDTA (TBE) buffer. Ten μL of each genomic DNA sample was loaded into each well and 1X TBE Buffer was used for electrophoresis to ensure adequate buffering power during vertical electrophoresis. The gels were electrophoresed at room temperature for ~60 min at 200 V (BLOT POWER-BP-T8, Biocraft, Japan) and stained with Midori Green (Nippon Genetics, Japan). The time of electrophoresis may need to be adjusted to the electrophoresis apparatus used. A DNA size marker (Gene Ruler Ultra Low Range DNA Ladder, Thermo Scientific, SM1211) was used. Images were acquired with FAS-Digi LED Imaging System (Nippon Genetics, Japan). Finally, the exact mutations in the selected screened samples were identified by direct Sanger sequencing. The list of primers used for Sanger sequencing is listed in Table 2.

**Table 2.**
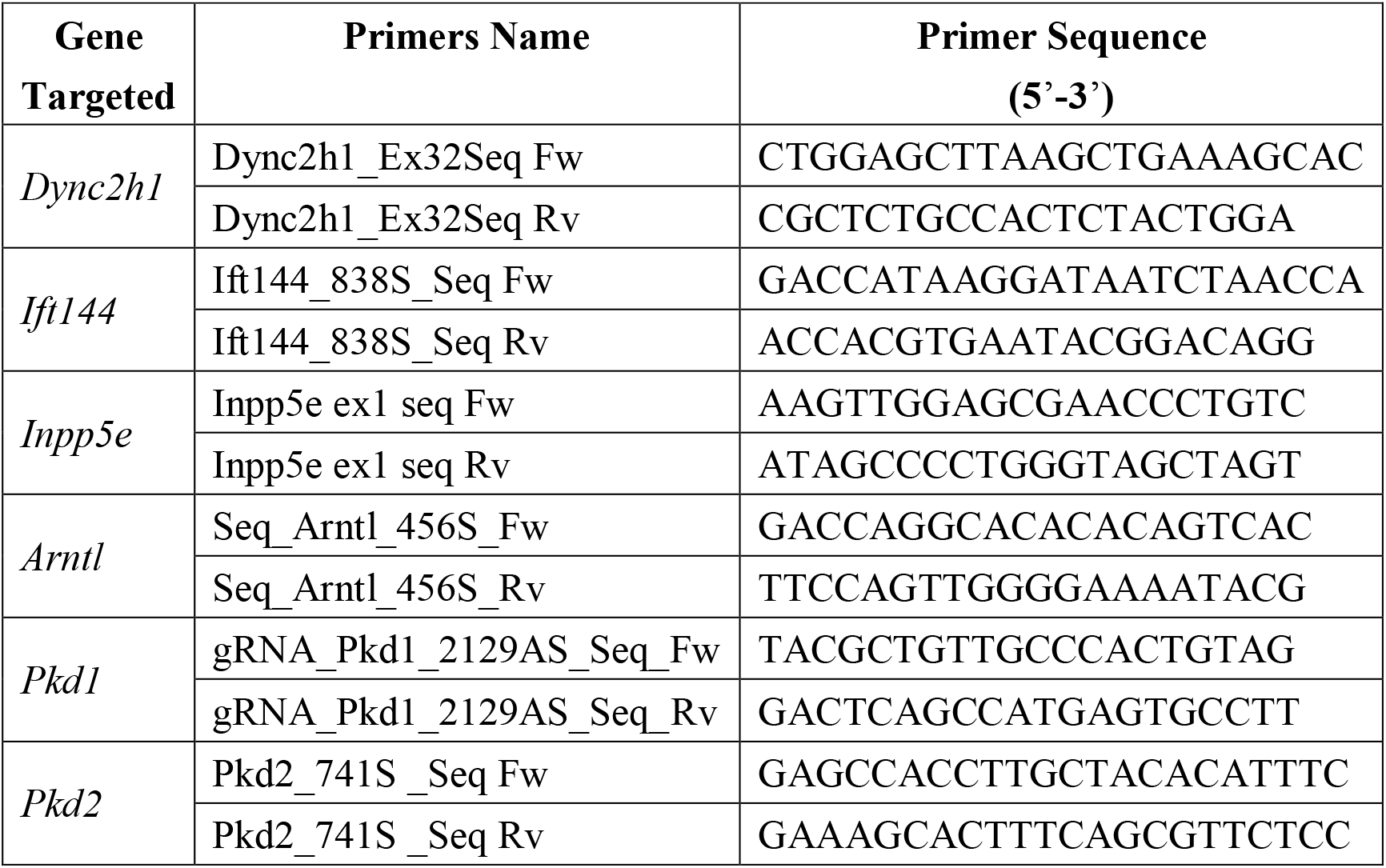
Sequencing primers.

### Antibodies

The antibodies used in this study are as follows: Arl13b (mouse mAb N295B/66; ab136648; Abcam), Arl13b (rabbit pAb; 17711-1-AP; Protein Tech), Alexa fluorophore-conjugated secondary antibodies (Thermo) for immunofluorescence microscopy.

### Immunocytochemistry (ICC)

Cultured cells were fixed with 4% paraformaldehyde (PFA, pH7.5) for 30 min at 37°C. Cells were washed with PBS and were blocked and permeabilized with 5% normal goat serum containing 0.1% Triton X-100 in PBS for 1 h at room temperature followed by incubation with the primary antibodies for 24 h at 4°C. After washing in PBS, cells were incubated for 2 h at room temperature with Alexa Fluor-conjugated t secondary antibodies and DAPI (1:1000; DOJINDO). Images were acquired using a confocal microscope (Olympus FV1000) equipped with an oil immersion lens (60X, NA 1.35).

## Results

### Concept and design of DST-PCR coupled with TBE-High-Resolution-PAGE

A key element of DST-PCR assay is the primers, which are sensitive to indels at the Cas9-mediated DSB. We designed “face-to-face” primers that annealed to sequences in the genomic DNA spanning the targeted Cas9 DSB site with both forward and reverse primers 3’ ends facing each other at the position between 3-bp and 4-bp upstream of the PAM sequence (Fig. 1a). The face-to-face primers generate a small amplicon ranging in size from 39-to 41-bp. After amplification with DST-PCR, PCR products are directly subjected to TBE-high-resolution PAGE, which contains high concentration of bisacrylamide, for mutant clones detection. The use of TBE-PAGE allows discriminating the size of indels with 1-bp resolution. Amplicons generated from mutant alleles that harbor insertion mutations will be detected as upper band shifts depending on the number of insertions (Fig. 1b). In contrast, target sequences with deletion mutations will not amplify because of the inability of the primers to anneal to the target sequences, especially the failure of 3’ end of primers resulting in no band on the gel (Fig. 1b). Similarly, bands will disappear when nucleotide substitution occurs at the site corresponding to the 3’ end of primers (Fig. 1b).

**Figure 1.**
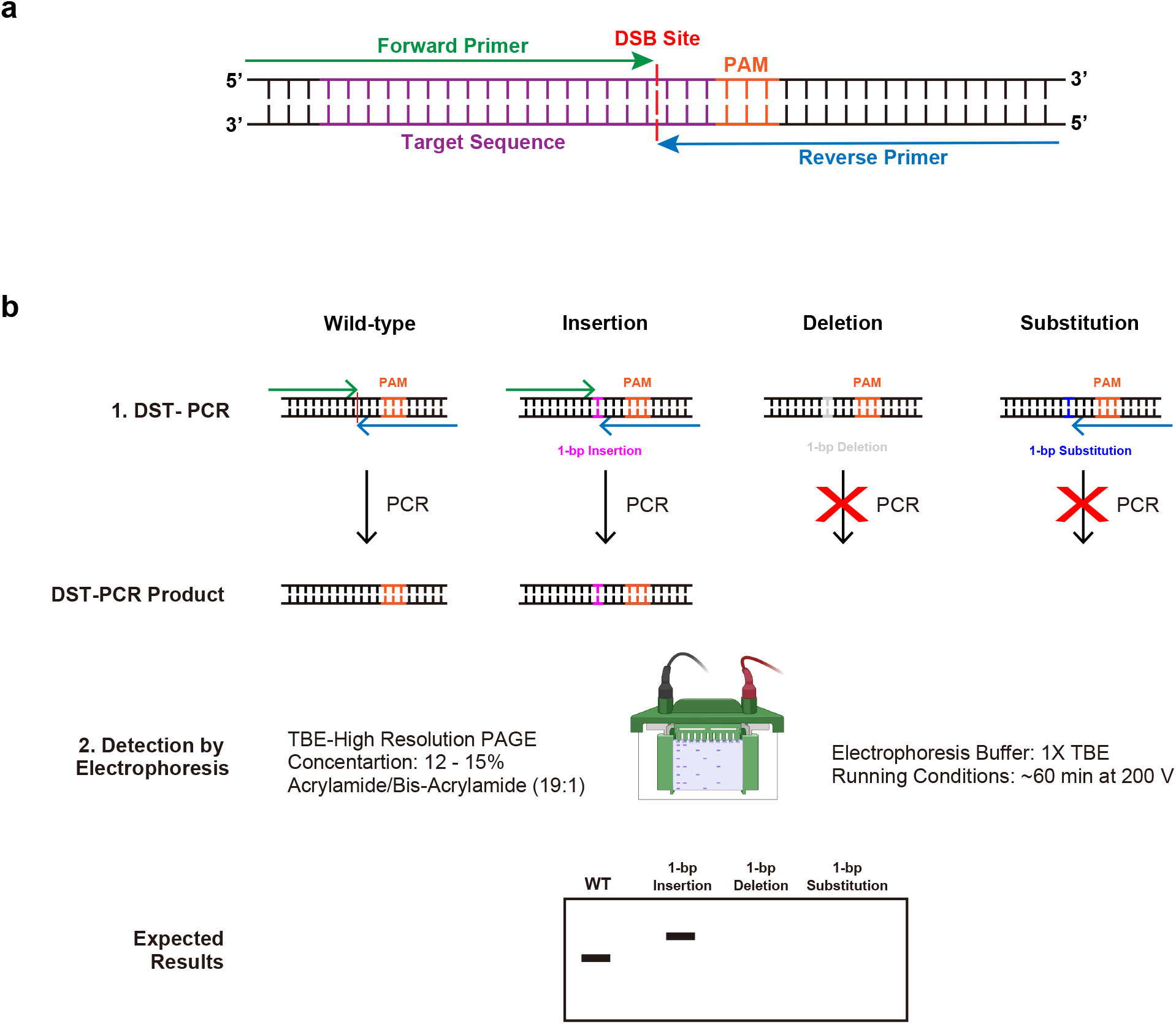
Overview of DST-PCR (<u>D</u>SB <u>S</u>ite-<u>T</u>argeted PCR) **(a)** Principle of DST-PCR. Target sequences, PAM sequences, and double-strand break (DSB) are shown in magenta, orange, and red, respectively. Forward primer (green arrow) and reverse primer (blue arrow) face each other at the DSB site, i.e. between 3-bp and 4-bp upstream of the PAM sequence. **(b)** Overview of DST-PCR. 1. Principle of DST-PCR in distinguishing indels and wildtype sequences. Black lines represent genomic DNA from an allele of WT, 1-bp insertion (pink) mutant, 1-bp deletion (grey) mutant, and 1-bp substitution (Navy). 2. Parameters for running TBE-PAGE and expected result of DST-PCR resolved by TBE-high-resolution-PAGE. Note that TBE-high-resolution-PAGE represents simplified pictures, it is possible that more than a single extra signal may be observed due to the nature of indels.

### DST-PCR coupled with TBE-High-Resolution-PAGE can efficiently screen insertion mutations induced by CRISPR/Cas9 gene editing

To evaluate the DST-PCR method, we first generated the CRISPR/Cas9-mediated NIH/3T3-*Dync2h1*-KO cells using sgRNA targeting exon 32 of murine *Dync2h1* gene locus (Fig. 2a). Dync2h1 is one of two cytoplasmic dynein heavy chain proteins that is responsible for the generation and maintenance of primary cilia [18]. Distinguishable band patterns between control cells and mutant clones were obtained from TBE-high-resolution-PAGE after DST-PCR. Control bands were detected at 40-bp, while clone #7 showed a 2-bp upper band shift (* of Fig. 2b), indicating insertion of 2 nucleotides into the target region of the *Dync2h1* gene on at least one allele. We next confirmed the insertion mutations found in the *Dync2h1*-KO clone #7 using Sanger sequencing and observed overlapping peaks in the sequencing chromatographs (Fig. 2c). Two nucleotides (AT) were inserted between −8 and −7 from the third nucleotide (G) of the PAM motif (AGG), hereafter called position 0, in one allele (Fig. 2c). In the other allele, a 1-bp deletion occurred at −7 from position 0 (Fig. 2c), which failed to be detected in PAGE (Fig. 2b). Both these mutations result in frameshift missense mutation leading to the heterozygous biallelic functional knockout of the *Dync2h1* gene. To confirm the loss of Dync2h1 protein in the selected clone, we examined whether the cells showed deficiency in primary cilia using an antibody against the primary cilium marker Arl13b. Compared to wild-type cells, cells of clone #7 showed fewer primary cilia with shortened length (Fig. 2d) confirming the generation of the *Dync2h1*-KO cell line.

**Figure 2.**
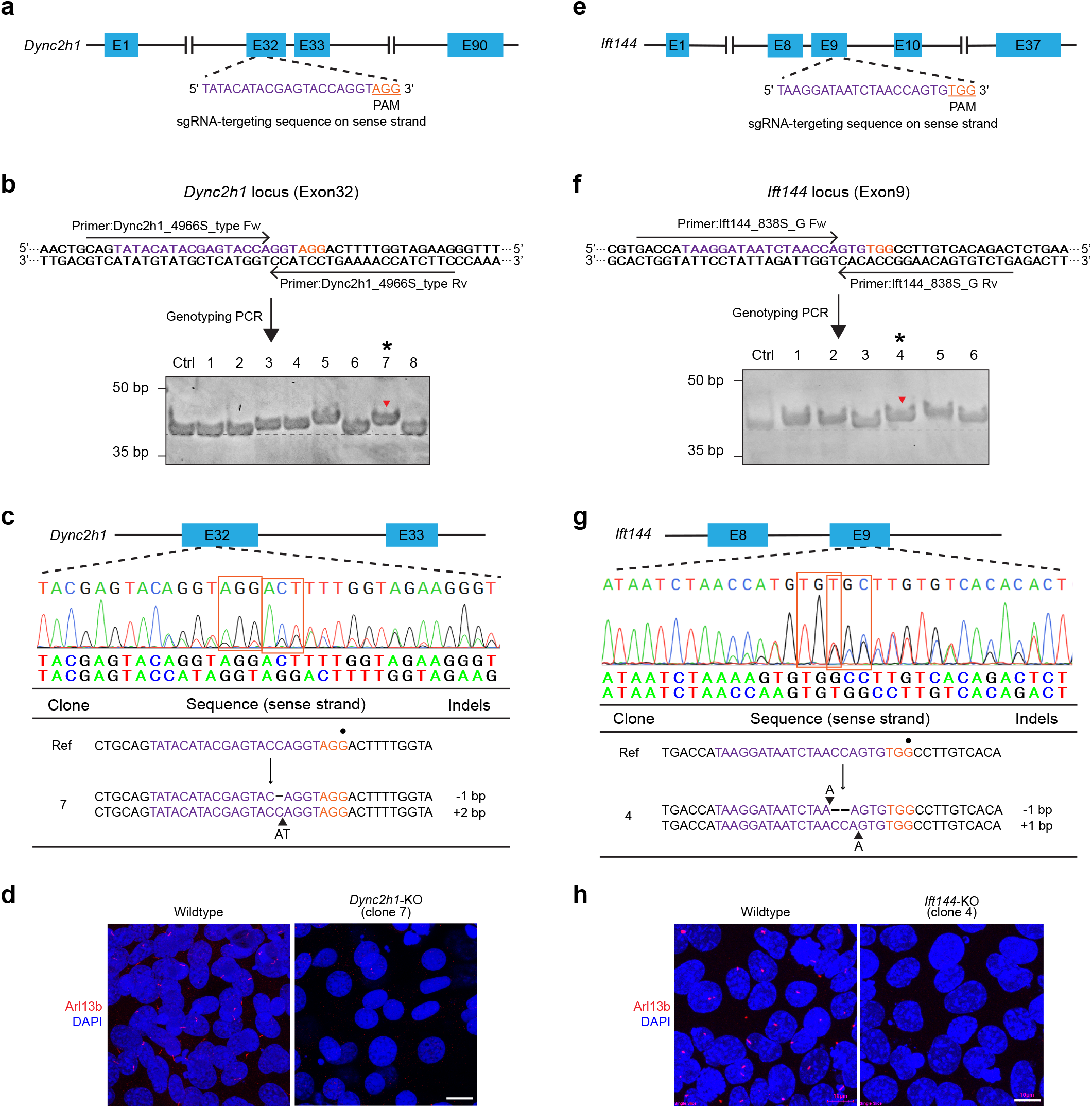
Detection of insertion mutations via DST-PCR. **(a)** Scheme of the Dync2h1 guide RNA targeting exon 32 in the murine *Dync2h1* gene. **(b)** DST-PCR for screening mutations. Upper panel: The arrows indicate the locations of the DST-PCR primers (see Table 1). Bottom panel: TBE-high-resolution-PAGE of DST-PCR products. Lane Ctrl: PCR product (40-bp) amplified from the genomic DNA of wildtype unedited cells. Several clones including clone #7 (*) show fragments mobility upper shift (red arrowhead). **(c)** Sanger sequencing of the PCR products of clone #7 (* in panel **b**) around the PAM sequence (orange rectangle). Split sequences were shown under the spectrum. **(d)** Fluorescence microscopy images showing cells with primary cilium stained for the ciliary marker (Arl13b; red) and the nucleus (DAPI; blue) in wild-type and clone #7. Scale bar, 10 μm. **(e)** Scheme of the Ift144 guide RNA targeting exon 9 in the murine *Ift144* gene. **(f)** DST-PCR for screening mutations. Upper panel: The arrows indicate the locations of the DST-PCR primers (see Table 1). Bottom panel: TBE-high-resolution-PAGE of DST-PCR products. Lane Ctrl: PCR product (41-bp) amplified from the genomic DNA of wildtype unedited cells. Several clones including clone #4 (*) show fragments mobility upper shift (red arrowhead). **(g)** Sanger sequencing of the PCR products of clone #4 (* in panel **f**) around the PAM sequence (orange rectangle). Split sequences were shown under the spectrum. **(h)** Fluorescence microscopy images showing cells with primary cilium stained for the ciliary marker (Arl13b; red) and the nucleus (DAPI; blue) in wild-type and clone #4. Scale bar, 10 μm. The 20-bp sgRNA and 3-bp PAM sequences are highlighted in magenta and orange respectively. Wild-type sequence (Ref), base deletion (-), base insertion (▲), position 0 of the PAM motif (●).

In addition, we applied DST-PCR to screen mutations in the mIMCD3 cells that have undergone CRISPR/Cas9 gene editing at exon 9 of endogenous murine intraflagellar transport protein 144 (*Ift144*) gene locus using Ift144 sgRNA to generate *Ift144*-KO clones (Fig. 2e). Genomic DNA was extracted from mutant cells derived from single-cell clones and subjected to DST-PCR. We found a 1-bp upper band shift in clone #4 (* of Fig. 2f) compared to control cells (41-bp band). Sanger sequencing of the clone# 4 showed that it was a heterozygous biallelic functional KO with a 1-bp (A) insertion (−6 from position 0) in one allele, and a 1-bp diminish by replacement of 2-bp nucleotides (CC) by a nucleotide A (−8 to −7 from position 0) in the other allele (Fig. 2g). Ift144 is a part of the IFT-A protein complex essential for anterograde transport in cilia and primary cilia formation [19–20]. To confirm the loss of Ift144 protein in the knockout cells, the primary cilium was visualized in clone #4 using an antibody against the primary cilium marker Arl13b. Compared to the wildtype cells, the clone #4 cells had no primary cilium (Fig. 2h), confirming the generation of *Ift144*-KO cell line and the feasibility of our screening approach to detect small insertion induced by CRISPR/Cas9 system with 1-bp resolution. These results suggest that DST-PCR along with TBE-high-resolution-PAGE is able to identify insertion mutations in cell clones engineered through CRISPR/Cas9 editing.

### DST-PCR coupled with TBE-High-Resolution-PAGE can detect deletion mutations induced by CRISPR/Cas9 gene editing

We further tested the efficiencies of DST-PCR along with TBE-high-resolution-PAGE to other genes. We attempted to generate KO mIMCD-3 cells of Inositol Polyphosphate-5-Phosphatase E (*Inpp5e*) gene that have undergone CRISPR/Cas9 gene editing using sgRNA sequences targeting exon 1 of murine *Inpp5e* locus as shown in Fig. 3a. Inpp5e compartmentalizes PIP in the primary cilium and regulates the primary cilium tip excision [15]. Target sequences were amplified using DST-PCR, and the amplicons were subjected to the TBE-high-resolution-PAGE, in which control bands were detected at 39-bp. Surprisingly, mutant clone #4 showed a 1-bp lower band shift at 38 bp as well as a 1-bp upper band shift at 40 bp (* of Fig. 3b). Sequencing results confirmed that clone 4 had a 1-bp deletion located at −5 from position 0 of the PAM motif (CCC) in one allele and a 1-bp (G) insertion (−6 from position 0) in the other allele (Fig. 3c).

**Figure 3.**
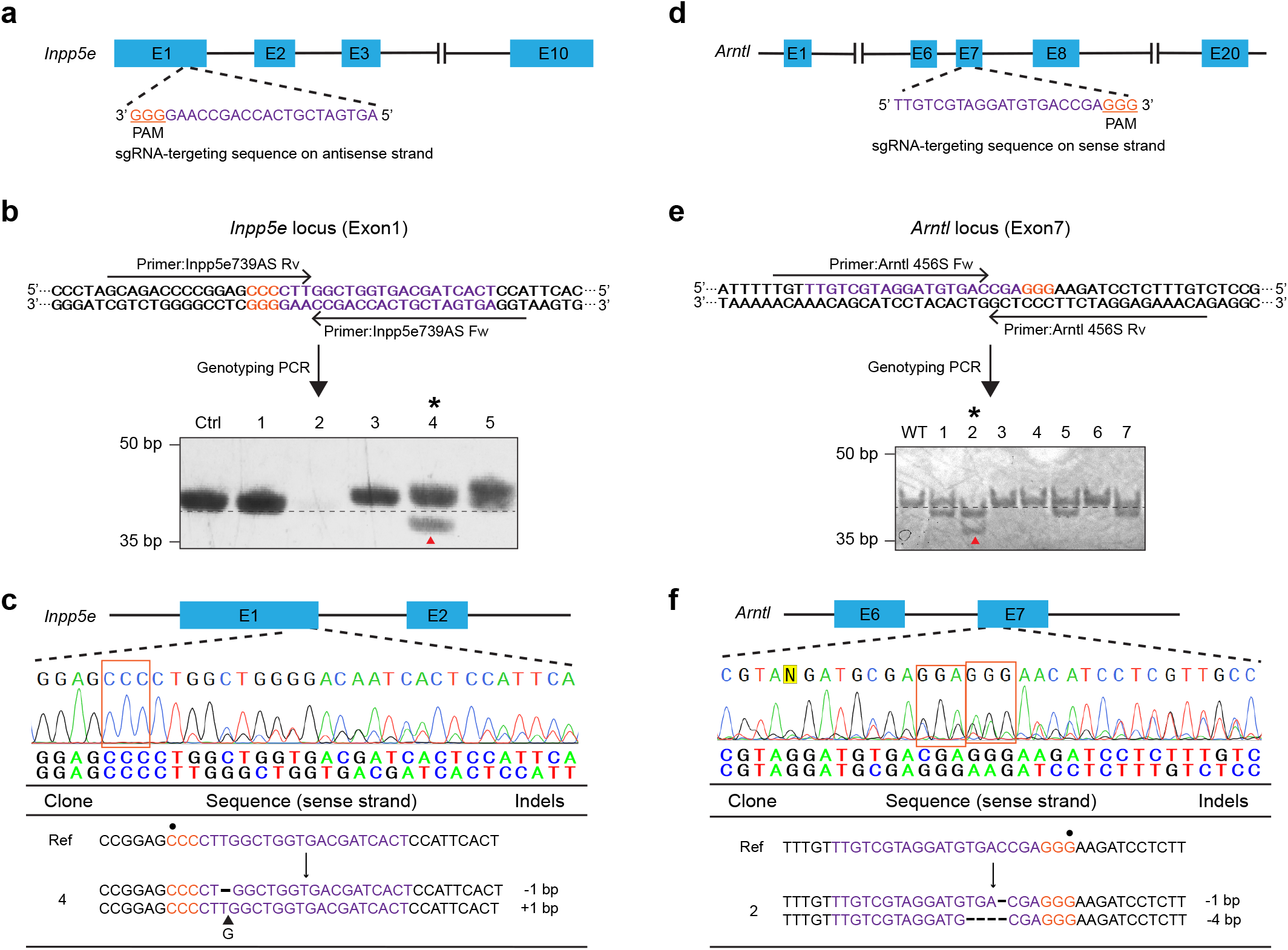
Detection of deletion mutations via DST-PCR. **(a)** Scheme of the Inpp5e guide RNA targeting exon 1 in the murine *Inpp5e* gene. **(b)** DST-PCR for screening mutations. Upper panel: The arrows indicate the locations of the DST-PCR primers (see Table 1). Bottom panel: TBE-high-resolution-PAGE of DST-PCR products. Lane Ctrl: PCR product (39-bp) amplified from the genomic DNA of wild-type unedited cells. Clone #4 (*) exhibited a lower shifted band (red arrowhead) **(c)** Sanger sequencing of the PCR products of clone #4 (* in panel **b**) around the PAM sequence (orange rectangle). Split sequences were shown under the spectrum. **(d)** Scheme of the Arnt1 guide RNA targeting exon 7 in the murine *Arnt1* gene. **(e)** DST-PCR for screening mutations. Upper panel: The arrows indicate the locations of the DST-PCR primers (see Table 1). Bottom panel: TBE-high-resolution-PAGE of DST-PCR products. Lane Ctrl: PCR product (40-bp) amplified from the genomic DNA of wild-type unedited cells. Clone #2 (*) exhibited two lower shifted bands (red arrow head). **(f)** Sanger sequencing of the PCR products of clone #2 (* in panel **e**) around the PAM sequence (orange rectangle). Split sequences were shown under the spectrum. The 20-bp sgRNA and 3-bp PAM sequences are highlighted in magenta and orange respectively. Wild-type sequence (Ref), base deletion (-), base insertion (▲), position 0 of the PAM motif (●).

We also analyzed NIH/3T3 cells clones lacking the aryl hydrocarbon receptor nuclear translocator-like (*Arntl*) gene. *Arntl* encodes Brain and muscle aryl hydrocarbon receptor nuclear translocator-like protein-1 (Bmal1) clock gene protein which is an essential component of the complex that regulates circadian rhythm in mammals [21]. We identified sgRNA sequences targeting exon 7 of the *Arntl* gene locus (Fig. 3d) and designed primers for screening as shown in Fig. 3e. After DST-PCR amplification, control bands were detected at 40-bp in the TBE-high-resolution-PAGE. Clone #2 exhibited lower band shifts of amplicons from both two alleles with a 1-bp and a 4-bp faster migration (* of Fig. 3e). Comparison of sequences near the *Arntl* sgRNA target site in clone #2 with the corresponding region in control cells showed that clone 2 had a 1-bp deletion located at −6 from position 0 of the PAM motif (GGG) in one allele and a 4-bp deletion (−9 to −6 from position 0) in the other allele (Fig. 3f). These findings demonstrate that DST-PCR is able to screen deletion mutations in cell clones engineered through CRISPR/Cas9 editing.

### DST-PCR coupled with TBE-High-Resolution-PAGE can efficiently screen homozygous biallelic indel mutations induced by CRISPR/Cas9 gene editing

CRISPR/Cas9 system allows the efficient production of homozygous biallelic KO cells [22]. To assess the potential of DST-PCR to detect homozygous indel mutations, we attempted to induce DSB in exon 11 of endogenous murine polycystic kidney disease 1 gene (*Pkd1*) after CRISPR/Cas9 gene editing using Pkd1 sgRNA in mIMCD-3 cells (Fig.4a). DST-PCR was performed using genomic DNA extracted from clones and control wild-type cells to amplify a 39-bp fragment corresponding to the target site in exon 11 of murine *Pkd1*. Clone 1 produced a single band with a 1-bp upper band shift compared to control cells (Fig. 4b). Sequencing results confirmed that clone #1 had a 1-bp homozygous biallelic insertion located at −6 from position 0 of the PAM motif (AGG) (Fig. 4c).

**Figure 4.**
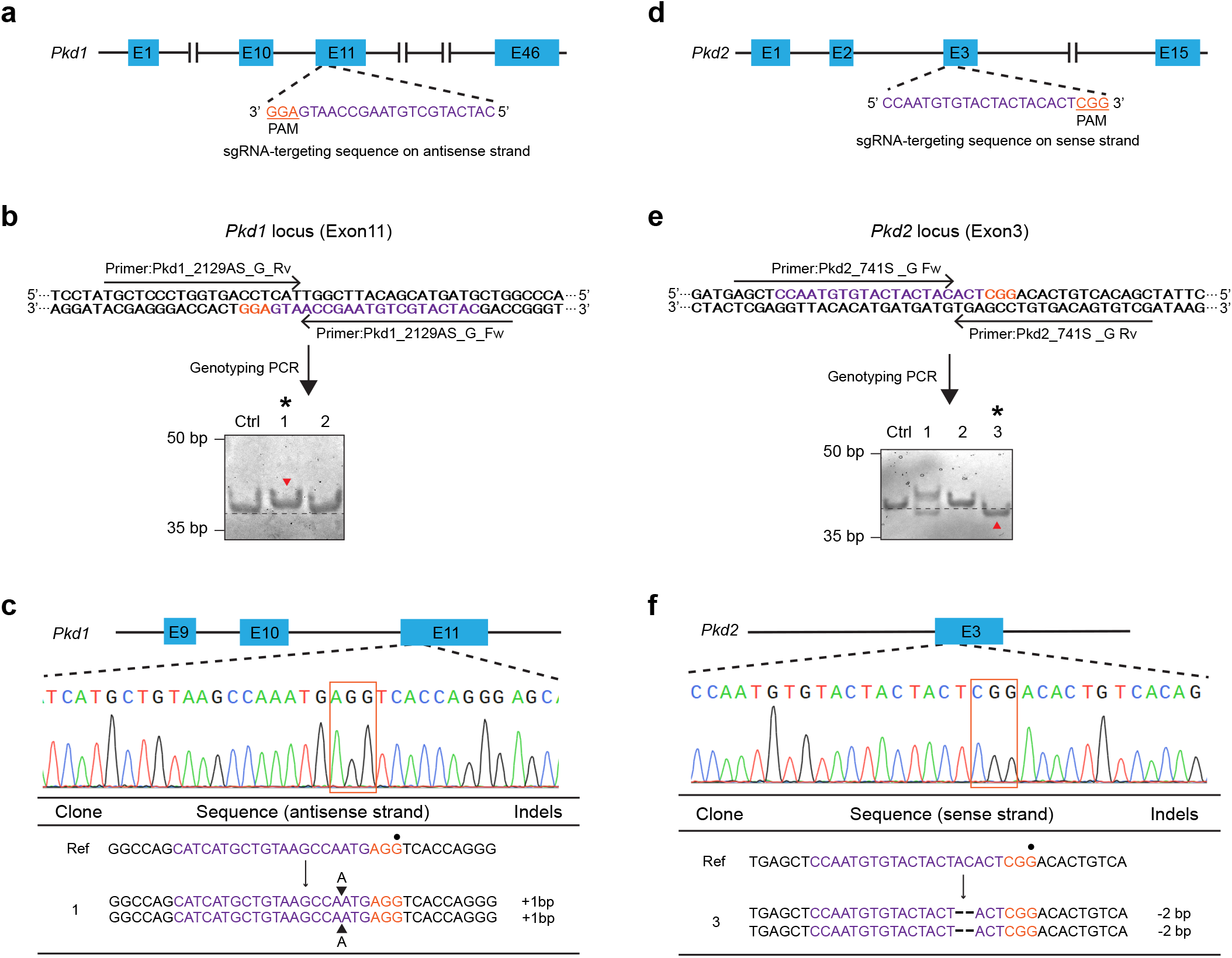
Detection of homozygous biallelic mutations via DST-PCR. **(a)** Scheme of the Pkd1 guide RNA targeting exon 11 in the murine *Pkd1* gene. **(b)** DST-PCR for screening mutations. Upper panel: The arrows indicate the locations of the DST-PCR primers (see Table 1). Bottom panel: TBE-high-resolution-PAGE of DST-PCR products. Lane Ctrl: PCR product (39-bp) amplified from the genomic DNA of wild-type unedited cells. Clone #1 (*) showed a fragment mobility upper shift (red arrowhead). **(c)** Sanger sequencing of the PCR products of clone #1 (* in panel **b**) around the PAM sequence (orange rectangle). **(d)** Scheme of the Pkd2 guide RNA targeting exon 3 in the murine *Pkd2* gene. **(e)** DST-PCR for screening mutations. Upper panel: The arrows indicate the locations of the DST-PCR primers (see Table 1). Bottom panel: TBE-high-resolution-PAGE of DST-PCR products derived. Lane Ctrl: PCR product (40-bp) amplified from the genomic DNA of wild-type unedited cells. Clone #3 (*) showed fragment mobility lower shift (red arrowhead). **(f)** Sanger sequencing of the PCR products of clone #3 (* in panel **e**) around the PAM sequence (orange rectangle). The 20-bp sgRNA and 3-bp PAM sequences are highlighted in magenta and orange respectively. Wild-type sequence (Ref), base deletion (-), base insertion (▲), position 0 of the PAM motif (●).

As another example, homozygous indel mutations in exon 3 of endogenous murine polycystic kidney disease 2 gene (*Pkd2*) were screened after CRISPR/Cas9 gene editing using Pkd2 sgRNA in mIMCD3 cells (Fig.4d). After DST-PCR, mutant clone #3 showed a 2-bp lower band shift (* of Fig. 4e) while the control band was detected at 40-bp (Fig. 4e). Comparison of indel sequences near the Pkd2 sgRNA target site in clone #3 with the corresponding region in control cells showed that clone #3 had a 2-bp homozygous deletion (−7 to −6 from position 0 of the PAM motif CGG) as shown in Fig. 4f. These data show that DST-PCR is able to screen homozygous biallelic indel mutations in cell clones engineered through CRISPR/Cas9 editing.

## Discussion

In this study, we described DST-PCR as a simple method to screen KO candidates generated by the CRISPR/Cas9 system before the final selection of clones with sequencing. To demonstrate our method, we implemented it to identify insertions and deletions in genetically altered NIH/3T3 and mIMCD3 cell clones generated from CRISPR/Cas9 gene editing. Indels in one or both alleles were detected using DST-PCR followed by 12-15% TBE-PAGE of the amplicons with a high concentration of bisacrylamide: 1-bp to 2-bp insertion mutations were detected in our hands. In the experiments for *Ift144* or *Dync2h1* KO, we detected amplicons only from one allele that harbors insertion mutations, while the homologous regions of the other alleles where a deletion or nucleotide substitution occurred were not amplified as initially expected (Figs. 2b and 2f). The failure of amplification of target region of the allele harboring deletion or substitution mutations could also be attributed to allelic drop out during PCR and also the fact that extension of the annealing failure is allele and mismatch position-dependent [23–25].

Interestingly, with this approach, we were able to detect 1-bp to 4-bp deletion mutations (Figs. 3b and 3e). The potential mechanism that enables DST-PCR to detect these types of mutations is that the mismatched nucleotide at the primer 3’ end was removed by the proofreading 3’ to 5’ exonuclease activity of the high-fidelity DNA polymerases [26–28] such as ExTaq DNA polymerase and KOD-Plus-Neo DNA polymerase during polymerizing and hence restored the PCR amplification capacity in part or completely. This means that DNA polymerases play an important role in deciding the discrimination ability of DST-PCR with high-fidelity DNA polymerase, increasing its ability in distinguishing the indels. Yet, a downside of using proofreading DNA polymerases is that it could introduce a bias towards PCR priming efficiency [29]. One way to overcome this problem and to detect deletion mutations more stably is to modify the screening primer pair design such that the 3’ ends of both forward and reverse primers are designed to face beyond 1-to 2-bp gaps on the DSB site. We applied this modified primer design and were able to enhance the screening power for deletion mutations (data not shown).

The detection of deletion mutations can also occur independent of the 3’ to 5’ exonuclease proofreading activity of the DNA polymerase at the primer 3’ end mismatch, which depends on the sequence around the DSB site. The 1-bp deletion of a nucleotide C in one of the alleles of the target sequence of the *Arntl* gene changed the sequence from 5’-GACCGA-3’ to 5’-GACGA-3’ still enabling the DST-PCR screening primer pair to bind to DNA templates as the C/G complementarity was kept for the 3’ end of the primers (Fig. 3e and 3f). Similarly, 2-bp deletion of nucleotides AC at the DSB site of the *Pkd2* gene was also detected (Fig. 4e). The sequence change from 5’-TACACT-3’ to 5’-TACT-3’ did not cause the mismatch between the primers and DNA templates (Fig. 4f). Both forward and reverse primers 3’ end (TAC of forward, AGT of reverse) could anneal to the DNA templates with perfect complementarity at the DSB site leading to the amplification of the fragment.

Our approach overcomes several weak points of widely used commercially available genotyping assays such as T7E1/Surveyor. These assays do not provide information for base number of indels as well as types of mutations. DST-PCR was able to detect the number of bases of indels with 1-bp resolution. Moreover, with the mismatch assays, a possibility of false-positive results remains [10, 30]; which is not an issue with the DST-PCR approach. These assays may lead to inaccurate conclusions as they are unable to distinguish between monoallelic mutants and heterozygous biallelic mutants [10, 31, 32], resulting in failure of excluding monoallelic mutants. Our approach can strongly screen out heterozygous monoallelic mutants that possess an unedited wildtype allele by excluding clones that show migration of bands identical to the wildtype sample. In addition, the mismatch assays technically miss homozygous biallelic mutants [10, 31, 32]. Our approach can detect homozygous biallelic mutants (Fig. 4).

The other benefit of our approach is the simplicity. Previously developed methods such as T7E1/Surveyor, RGEN-RFLP, DNA melting analysis [33], and fluorescent PCR [31] require additional enzymes, denaturing and hybridization of PCR products, specific oligonucleotide probes, and/or special chemicals. Our DST-PCR approach is truly a simple PCR which does require neither additional steps, special enzymes, nor chemicals. As only positive clones screened through DST-PCR need to be confirmed, it could lower the cost for subsequent confirmation by Sanger sequencing.

Direct sequencing of PCR products without the need for cloning individual PCR fragments is compatible with our approach. Except the mutation is homozygous, direct Sanger sequencing of DNA from diploid organisms with heterozygous mutations generates two overlapping traces at the mutation start site. When clones screened through DST-PCR and TBE-high-resolution PAGE are selected for Sanger sequencing, the information obtained about base numbers of indels is also helpful for subsequent sequence data analysis. With this prior knowledge of where and how many nucleotides are inserted or deleted in at least one allele, two overlapping traces can be easily interpreted.

Our approach still has some limitations to our approach though. The DST-PCR can only detect mutations close to the PAM, which are most frequent when using the CRISPR/Cas9 system, and cannot be used to detect mutations far from the PAM sequence. In addition, as our approach is based on PCR, there is a possibility of allele drop-out [23–25] and the PCR step may be biased towards the more abundant template and a small number of mutated sequence may not be detected [29, 34], for example in case of compound heterozygotes where one mutated template is more abundant than the other mutated template. Therefore, it is necessary to sequence the target region to confirm the outcome of genome editing. Furthermore, there may be limitations to designing primers for genotyping in high GC content regions or highly repetitive sequences in the target region [35–37]. Finally, there might be nonspecific signals that are shared among the lanes in WT and mutant clones. However, this problem can be solved by carefully designing and optimizing the PCR stringency conditions to avoid non-specific bands.

In summary, we have established a simple, purely usual PCR-based genotyping DST-PCR method that does not require any additional equipment, probes, nor special techniques to screen for indels in cell lines generated by CRISPR/Cas9 system. Our data demonstrate that this approach can identify both on-target insertions and deletions irrespective of the zygosity of the mutants in cell lines, supporting the general applicability and reproducibility of DST-PCR. It will allow researchers to screen for mutant clones without the need for special enzymes or devices in the lab and can be implemented with standard molecular biology equipment. While we did not yet try to genotype transgenic cell lines generated by zinc-finger nucleases (ZFN) [38] and transcription activator-like nuclease (TALEN) [39], this method will likely be applicable, as DNA modifications through NHEJ with these systems is fundamentally the same as with the CRISPR/Cas9 system.

## Acknowledgments

This work was supported in part by Grants-in-Aid for Scientific Research Activity Startup 19K23728 and for Early-Career Scientists 21K15088 (to F.I.), a Grant-in-Aid for Early-Career Scientists 20K15792 (to R.N.), and Grant-in-Aids for Scientific Research on Innovative Areas 15H01207 and for Scientific Research (C) 21K06172 and Japan Science and Technology Agency, Precursory Research for Embryonic Science and Technology JPMJPR17H1 (to K.I.) We thank Madoka Hamada, Mimoko Katoh, and lab members for their technical assistance and useful discussions.

## Author contributions

K.I. designed research; F.I., R.N., and K.I. performed experiments and acquired the data; F.I., R.N., M.S., and K.I. analyzed the data; F.I., R.N., and K.I. wrote the paper; all authors discussed the results and commented on the manuscript.

## Data Availability

All data generated or analysed during this study are included in this published article.

## Additional Information

Competing Interests: The authors declare no competing interests.

